# Optimal inter-electrode distances for maximizing single unit yield per electrode in neural recordings

**DOI:** 10.1101/2025.04.11.648355

**Authors:** Domokos Meszéna, Ward Fadel, Róbert Tóth, Angelique C. Paulk, Sydney S. Cash, Ziv Williams, Tamás Kiss, Marcell Stippinger, Lucia Wittner, Richárd Fiáth, Zoltán Somogyvári

## Abstract

State-of-the-art high-density multielectrode arrays enable the recording of simultaneous spiking activity from hundreds of neurons. Although significant efforts have been dedicated to enhancing neural recording devices and developing more efficient sorting algorithms, there has been relatively less focus on the allocation of microelectrodes—a factor that undeniably affects spike sorting effectiveness and ultimately the total number of detected neurons. Here, we systematically examined the relationship between optimal electrode spacing and spike sorting efficiency by creating virtual sparser layouts from high-density recordings through spatial downsampling. We assessed spike sorting performance by comparing the quantity of well-isolated single units per electrode in sparse configurations across various brain regions (neocortex and thalamus) and species (rat, mouse, and human). Enabling the theoretical estimation of optimal electrode arrangements, we complement experimental results with a geometrical modeling framework. Contrary to the general assumption that higher electrode density inherently leads to more efficient sorting, both our theoretical and experimental results reveal a clear optimum for electrode spacing specific to species and regions. We demonstrate that carefully choosing optimal electrode distances could yield a total of 1.7 to 3 times increase in spike sorting efficiency. These findings emphasize the necessity of species- and region-specific microelectrode design optimization.

## Introduction

Extracellular multielectrode recordings and spike sorting are among the most essential techniques for investigating neural circuits. Modern high-density, silicon-based probes contain thousands of closely packed microelectrodes, allowing the capture of the simultaneous activity of hundreds of neurons with high spatiotemporal resolution^1–4^. Since these complementary metal-oxide semiconductor (CMOS)-based devices typically have fewer recording channels than physical microelectrodes, only a small subset of electrodes can be used for simultaneous recordings at a time. Thus, sequential mapping of neuronal activity across all electrodes within the brain tissue further increases the neuron yield, reaching thousands of individual neurons that can be monitored in a single subject^3^. Despite these impressive unit yields, the average number of neurons detected per a single electrode remains relatively low^5,6^, while the vast amounts of data generated in these experiments pose significant challenges for storage and processing^7^.

Such high-density silicon probes (i.e., Neuropixels probes) have also recently been applied in human studies^8,9^, suggesting that it is likely just a matter of years before high-density, high-channel-count brain implants find their way into clinical brain-computer interface (BCI) applications and sensory neuroprostheses^10–12^. However, despite substantial progress, wireless invasive BCI devices still suffer from a bandwidth bottleneck. Efficient transmission and processing of neural data sampled at high rates (>10 kHz) from a large number of electrodes remains an unresolved issue, especially for real-time applications.

A potential solution to these challenges could be to strategically select a subset of microelectrodes to maximize the neuron yield per electrode. This approach could reduce the bandwidth required for BCI applications and decrease the amount of data generated in both acute and chronic animal experiments^13^. However, an optimal selection of electrodes based, for example, on adaptive strategies is typically computationally intensive, time-consuming, and requires regular updates due to changes in neural signals over time^13^. Another method involves pooling electrodes that contain high-amplitude spikes, but this technique demands considerable modifications to recording hardware and software and has yet to be demonstrated in practice^14^. In this study, we propose an alternative strategy: selecting electrodes at a distance optimized to maximize neuron detection per single electrode. This approach would be less resource-intensive, as users could simply select the adequate electrode arrangement from a list of predefined configurations with equidistant spacing between electrodes. However, for this method to be viable, the optimal inter-electrode distances must first be experimentally determined for different brain regions and species.

Currently, hundreds of different electrode configurations are used for neural recordings. Based on the few experimental studies that have examined the effect of electrode configuration on the yield of well-isolated neurons, we know that the spatial arrangement of microelectrodes influences both the number of single units detected and the quality of spike sorting results^4,15,16^. These studies generally agree that the higher channel count (i.e., denser spatial sampling) increases the total neuron yield but have not assessed the efficiency of different electrode configurations in terms of neuron count per electrode. Recently, a modeling study compared different electrode configurations with a fixed electrode count to evaluate the single unit yield^17^. While the vertical spacing was fixed at 20*μm*, the two columns of 30 electrodes were horizontally spaced 16, 32 or 48*μm* apart or formed a staggered pattern. The findings revealed that the unit yield was the highest in the case of 48*μm* horizontal spacing, suggesting that a higher electrode density does not always lead to a higher unit yield.

Interestingly, the first study to establish a simple theoretical model linking electrode configurations to electrode efficiency was published only in 2021^18^. This model, known as the “dual observer model”, allows the calculation of optimal electrode configurations and inter-electrode distances for various recording conditions. In that work, the model and the model parameters were determined using simulated recordings, where real spike patterns were embedded into spectrum-fitted pink noise backgrounds at random positions. A key advantage of this simulation-based approach is that the ground truth is known, allowing for a precise evaluation of spike sorting performance. However, this approach has certain limitations. Most importantly, it does not consider the natural variability of spike waveforms or their non-stationarity during the recordings. Furthermore, it does not account for correlations between spikes and between the background noise and spikes. These factors may impact the spike sorting performance in ways that remain poorly understood. Thus, spike sorting in real in vivo extracellular measurements is likely more challenging than in simulated recordings.

In this study, we take a complementary approach by using in vivo multichannel recordings. The advantage of this real-data-based approach is that spike sorting algorithms are challenged by all the imperfections and complexities inherent in neuronal recordings. However, the lack of ground truth makes evaluating spike sorting performance in real data more challenging. Rather than using a ground-truth-based evaluation, here we rely on traditional quality metrics of single unit clusters, which are widely used in everyday laboratory practice. Thus, the aims of this study are the following:

- To assess spike sorting efficacy for different inter-electrode distances across various brain areas (neocortex and thalamus), species (rat, mouse and human), and spike sorting algorithms (Kilosort1 and Kilosort2).
- To test the validity of the previously introduced dual observer model for real in vivo neuronal data and different spike sorting algorithms.
- To determine and compare model parameters across different brain areas and species.
- To determine optimal electrode configurations for different brain areas and species.

### The dual observer model

First, we briefly summarize the dual observer model introduced by Tóth et al.^18^, which describes the factors that link electrode configurations to spike sorting efficiency. Numerous simulated spike sorting experiments have led us to this simplified description, which assumes that the spike sorting efficiency of an electrode configuration can be determined by three main factors.

The first factor is the volume of the neural tissue, *V*_*single*_, within which neurons are close enough to reach the signal-to-noise ratio threshold required for successful spike sorting. Here, we assume that this volume can be well approximated by spheres of radius *R* centered around the microelectrodes. We refer to *R* as the observation distance of an electrode (Fig. 1a).

**Figure 1.**
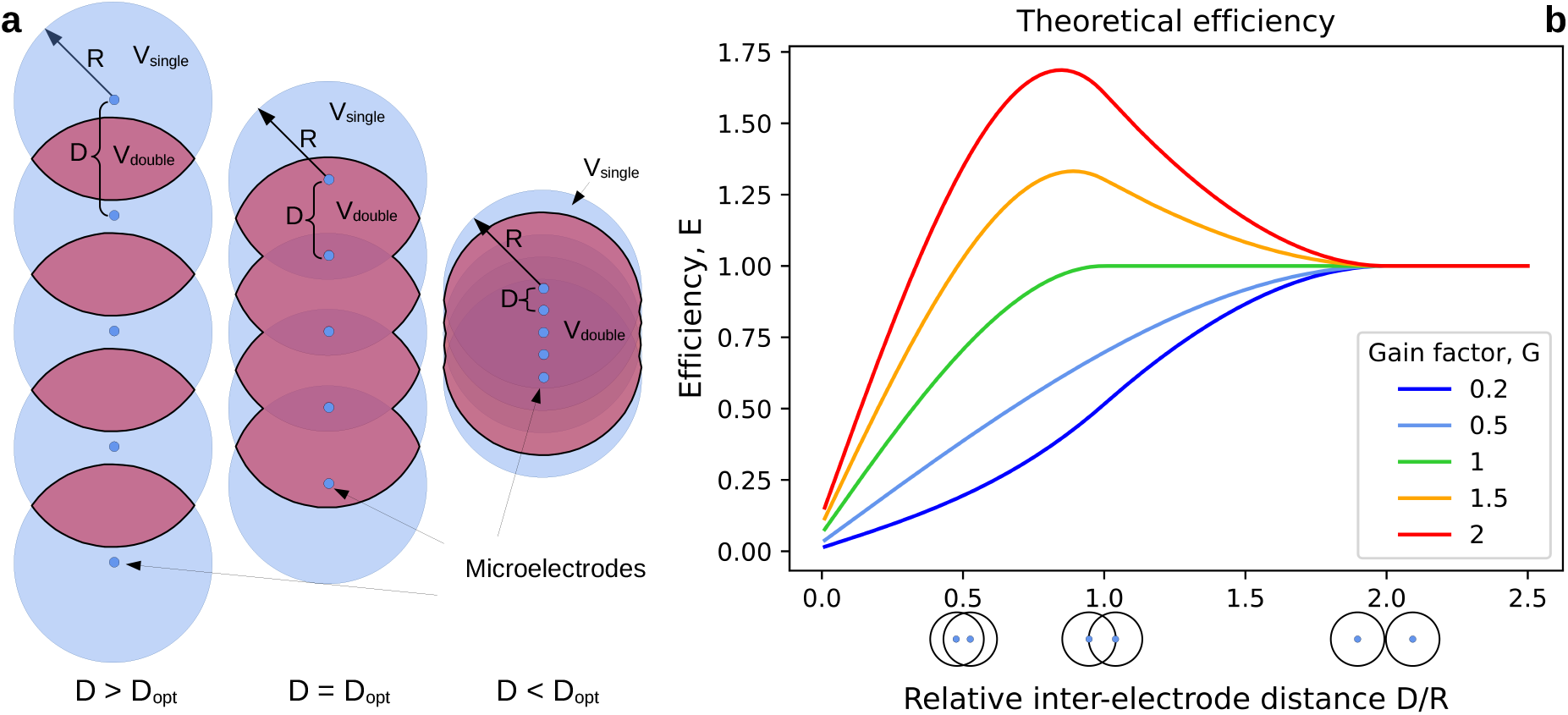
The dual observer model. (**a**) The circles around the microelectrodes (blue dots) represent the spheres (with an observation distance of R) within which spikes meet the signal-to-noise ratio threshold required for successful spike sorting. Thus, spikes are found within the blue volume with probability *p*_*single*_. The red region indicates the overlapping volume *V*_*double*_, where the probability of spike detection, *p*_*double*_, depends on the gain factor (*G*). The three cases illustrate different inter-electrode distances (*D*), where D is larger than (left), equal to (middle), or smaller than (right) the optimal distance *D*_*opt*_. **(b)** Spike sorting efficiency (*E*) as a function of relative inter-electrode distance, expressed in *D/R* units. The small pictograms below the *x*-axis illustrate the relative positions of observation sphere pairs at the corresponding *D/R* ratios: for *D/R* ≥ 2, there is no intersection; for 1 *< D/R <* 2, only first neighbors intersect; and for *D/R <* 1, intersections occur between farther neighbors as well. Efficiency is measured relative to independent electrodes, meaning that *E* converges to 1 for *D >* 2*R*. For *G <* 0.5 (blue), electrodes counteract each other. For 0.5 ≤ *G* ≤ 1 (light blue and green), electrodes positively cooperate, but the increased detection probability does not fully compensate for the loss of volume in the intersection. For *G >* 1 (orange and red), the gain from cooperation overrides the volume loss, resulting in a peak in efficiency at the optimal inter-electrode distance *D*_*opt*_.

The second factor is the probability of detecting a spike that can be reliably isolated within the observation distance, denoted as *p*_*single*_. This probability depends on the size and number of active neurons in a given volume, as well as the amplitude of their extracellular spikes and the noise level.

The third factor is the cooperation or synergy between electrodes, which quantifies how much more likely it is to detect a well-isolated spike in the intersecting volumes of the observation spheres compared to a volume observed by only a single electrode. We denote the volume of this double coverage as *V*_*double*_, which represents the tissue volume where spike amplitudes reach the detection threshold on at least two electrodes (Fig. 1a).

Thus, the number of observed high-quality single units, *N*, can be described as:

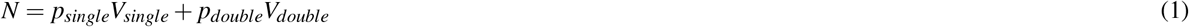

where *p*_*single*_ represents the probability density (probability per volume) of successful clustering based on single channel data, and *p*_*double*_ denotes the probability density of obtaining a high-quality single-unit cluster using two channels. Here, we assume that these two probabilities are disjoint.

By introducing the gain factor:

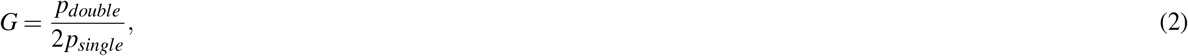

the model can be rewritten as:

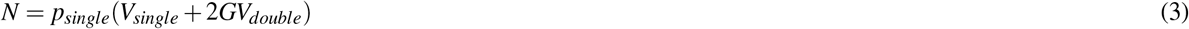

The gain factor *G* characterizes the type and efficiency of the interaction between electrodes. A value of *G <* 0.5 indicates negative cooperation between the electrodes, which means that the appearance of a spike on another electrode makes spike identification more difficult. A value of *G* = 0.5 signifies no cooperation between the electrodes—the sorting probability remains the same regardless of whether a neuron is observed on one channel or two.

For 0.5 *< G <* 1, there is a cooperative effect between the electrodes, but the increased observation probability in the overlapping regions does not fully compensate for the loss of total observed volume due to the intersections of the observation spheres.

At *G* = 1, the probability of detecting a high-quality unit in the overlapping regions is twice that of non-overlapping regions, fully compensating for the volume loss. In this case, the spike sorting efficiency does not depend on the inter-electrode distance (*D*) as long as *D > R*.

Finally, if *G >* 1, cooperation outweighs the volume loss, leading to an optimal inter-electrode distance *D*_*opt*_ that results in the highest efficiency.

To calculate *V*_*single*_ and *V*_*double*_, we need to know the electrode configuration, including the inter-electrode distance *D* and the observation distance *R*. Additionally, estimating the neuron yield requires knowledge of *p*_*single*_ and the gain factor *G*. Thus, given a specific electrode configuration, the model has three free parameters: the observation distance *R*, the spike density *p*_*single*_, and the gain factor *G*. These parameters are typically unknown and may depend on the brain area and the species being studied, but they can be determined by fitting the model to the experimental data.

As shown in Tóth et al.^18^, the dual observer model enables the calculation of optimal inter-electrode configurations and distances. Considering all regular one- and two-dimensional electrode arrangements, it was demonstrated that the theoretical optimum for a large number of electrodes is a hexagonal arrangement. However, one-dimensional linear electrode arrangements can achieve nearly the same efficiency if the inter-electrode distance is optimized.

Since one-dimensional (e.g., single-shank linear) probes are the most widely used electrode configurations, we focus exclusively on these linear probes and aim to determine their optimal inter-electrode distances. In addition to their practicality and high efficiency, linear electrode arrangements offer another advantage: they allow for an analytical calculation of spike sorting efficiency for any inter-electrode distance, given the model parameters (see Methods), as well as the determination of the optimal inter-electrode distance (Fig. 1b). Note that most of our results automatically generalize to multi-shank probes, provided that the inter-shank distance is greater than 2*R*.

For an array of *M >* 3 electrodes with even spacing, and given a known observation distance *R* and gain factor *G*, the optimal inter-electrode distance *D*_*opt*_ can be calculated as:

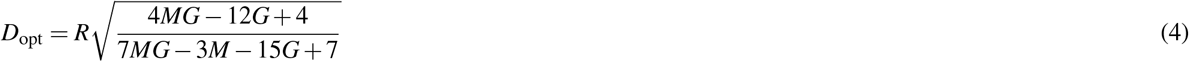

For *G* ≈ 1, this formula simplifies to *D*_*opt*_ ≈ *R*, while for *G* ≫ 1, the optimal inter-electrode distance converges to 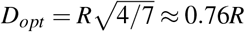.

Thus, the most important factor in determining the optimal inter-electrode distance is the observation radius *R*, as *D*_*opt*_ scales linearly with *R*. The second most influential factor is the gain factor *G*, which adjusts *D*_*opt*_ between *R* and 0.76*R*. On the other hand, *G* determines the peak efficiency achievable with the optimal electrode arrangement, as given by *E*_*opt*_ ≈ 0.76*G* + 0.16 (for calculation details, see Tóth et al.^18^).

Although the third parameter of the dual observer model, *p*_*single*_, is necessary for precise model fitting, it does not influence the optimal electrode spacing. Instead, it scales the unit-per-channel yield. These three parameters may vary across species, brain areas, and sorting algorithms. To explore these effects and parameters, we adopted the following approach in this study:

- First, we collected neuronal data obtained using high-density silicon probes in both rodents and human patients.
- Second, we systematically subsampled these high-density neuronal recordings and performed spike sorting on all measurements.
- Third, we calculated the median yield of well-isolated neurons per channel across all subsampled datasets.
- Fourth, we determined the model parameters *R, G*, and *p*_*single*_ for different brain areas and species by numerically fitting the model predictions to spike sorting results obtained from varying channel counts.
- Finally, we estimated the optimal inter-electrode distances for different brain areas and species.

## Results

To determine the parameters of the dual observer model and the optimal inter-electrode distances for different species (rat, mouse, and human) and brain areas (neocortex and thalamus), we first collected high-density neuronal data. For rodents, neuronal recordings were obtained using a 256-channel single-shank silicon probe from the neocortex and thalamus of rats (Fig. 2a,b; Supplementary Fig. 1a,b; Supplementary Table 1) and from the neocortex of mice (Fig. 2c; Supplementary Fig. 1c, Supplementary Table 2). The anatomical location of these recordings was identified based on the fluorescent track of the probe and Nissl-stained brain sections (Fig. 2e; Supplementary Tables 1-2). Additionally, we used a publicly available human dataset obtained with Neuropixels probes from the neocortex of patients undergoing deep brain stimulation surgery (Fig. 2d)^8,19^. The main properties of the rodent and human datasets are summarized in Table 1.

**Table 1.** Main properties of the four high-density neuronal datasets.

| Subject | Brain area | No. of recordings | No. of channels | Mean recording duration (min) |
| --- | --- | --- | --- | --- |
| Rat | Neocortex | 6 | 256 | 22.14 ± 3.87 |
| Rat | Thalamus | 5 | 256 | 16.98 ± 2.74 |
| Mouse | Neocortex | 7 | 256 | 24.18 ± 7.68 |
| Human | Neocortex | 2 | 384 | 14.25 ± 0.49 |

**Figure 2.**
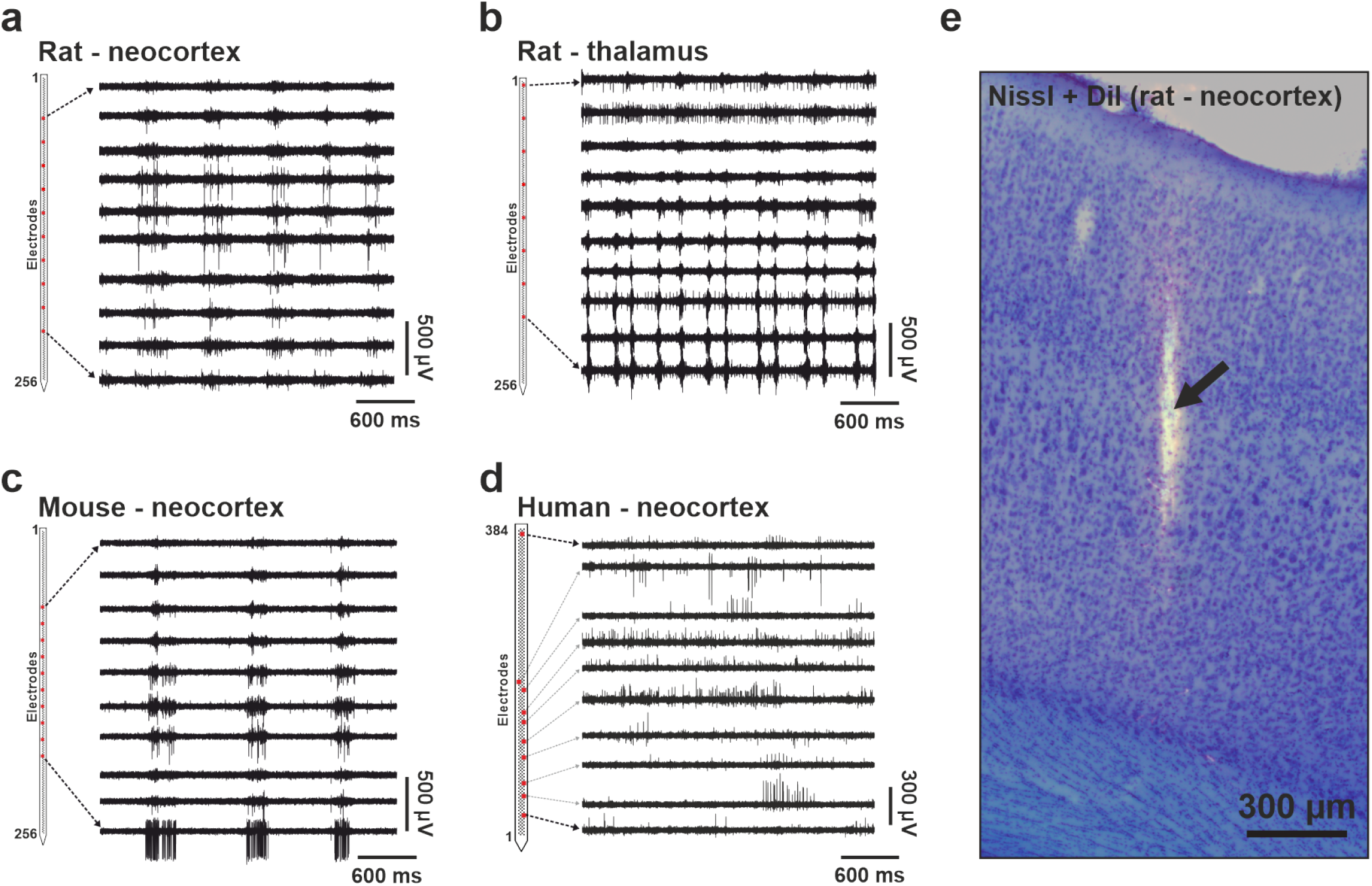
Representative 3-second-long examples of high-density neuronal recordings (500–5000 Hz frequency band) obtained from the rat neocortex (**a**), rat thalamus (**b**), mouse neocortex (**c**), and human neocortex (**d**). The rodent neural data was recorded with a custom 256-channel laminar silicon probe^15^, while the human data was acquired with a Neuropixels probe^8,19^. On the left, the schematic of the section of the probe shank containing the microelectrodes (small black squares) is displayed. For each species and brain region, spiking activity recorded on ten electrodes/channels (red squares) is shown. Rat and mouse data was acquired under ketamine/xylazine anesthesia, whereas the human patient (Pt02) was awake during recording. (**e**) A Nissl-stained coronal brain section showing a DiI-labeled probe track (black arrow) in the neocortex of a rat.

Next, for each dataset, we generated spatially downsampled, lower channel count recordings from the original, full-resolution data to emulate recordings with varying inter-electrode distances (Fig. 3; Table 2). These subsampled recordings had the same vertical coverage as the original recordings but lower spatial resolutions. Since only two human high-density recordings were available that had an appropriate quality for our analyses, we generated more recordings by using every possible electrode configuration with an equidistant spacing for a given channel count (i.e., inter-electrode distance; see Supplementary Fig. 2 for details). Then, we performed spike sorting both on the full-resolution and the spatially subsampled recordings to extract single units. To indicate the quality of unit isolation, in Fig. 4, we demonstrate multichannel spike waveforms of representative single units from each dataset, as well as the amplitude and depth distributions of isolated neurons. The final single unit yields are indicated in Fig. 5a and in Supplementary Tables 3-6. Additionally, for the rodent datasets, the distributions of computed quality metrics (see Methods for details) are shown in Supplementary Figure 3.

**Table 2.**
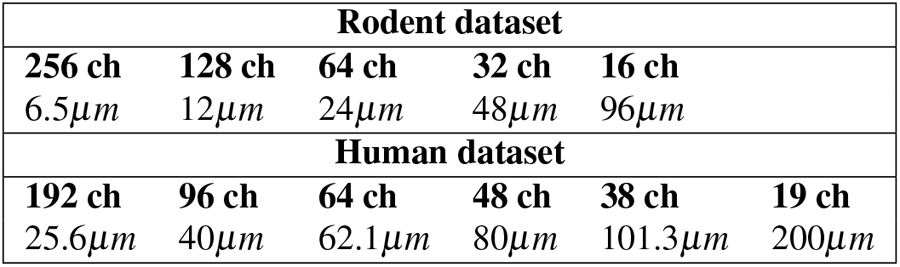
Inter-electrode distances for the original and the spatially downsampled recordings of the rodent and human datasets. Spatial resolution decreases from left to right. ch, channel.

| Rodent dataset |  |  |  |  |  |
| --- | --- | --- | --- | --- | --- |
| 256 ch | 128 ch | 64 ch | 32 ch | 16 ch |  |
| 6.5μm | 12μm | 24μm | 48μm | 96μm | |
| Human dataset |  |  |  |  |  |
| 192 ch | 96 ch | 64 ch | 48 ch | 38 ch | 19 ch |
| 25.6μm | 40μm | 62.1μm | 80μm | 101.3μm | 200μm |

**Figure 3.**
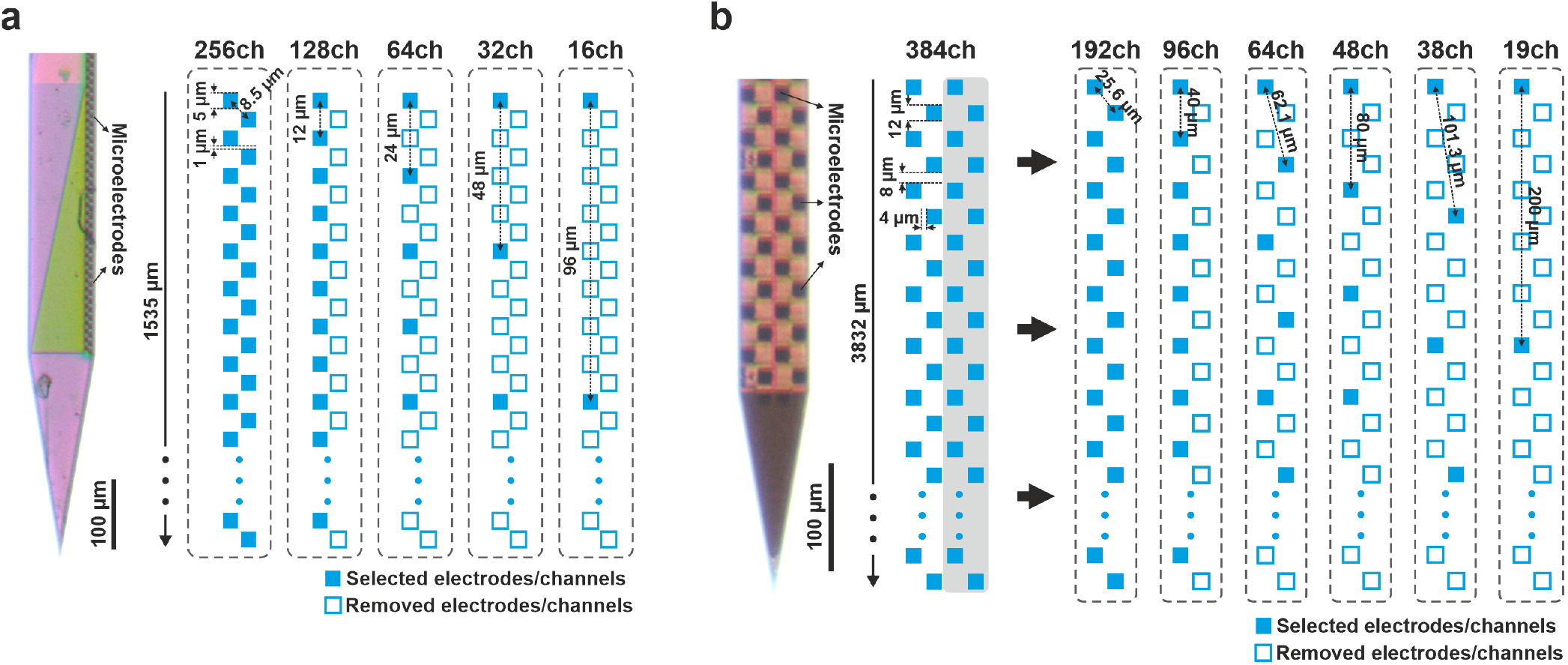
Spatial downsampling of high-density neuronal recordings obtained with 256-channel laminar silicon probes in rodents (**a**) and with Neuropixels silicon probes in humans (**b**). Lower channel (ch) count recordings, corresponding to reduced spatial resolutions, were generated from the original recordings by removing subsets of electrodes. The spatial resolution of recordings decreases from left to right. The size of the microelectrodes, along with the inter-electrode distances for both the original and subsampled recordings, is displayed. A stereomicroscopic image of the tip region of the silicon probes is shown on the left. For the Neuropixels recordings, only data recorded by the rightmost two columns of electrodes (gray shaded area) were used for analysis.

**Figure 4.**
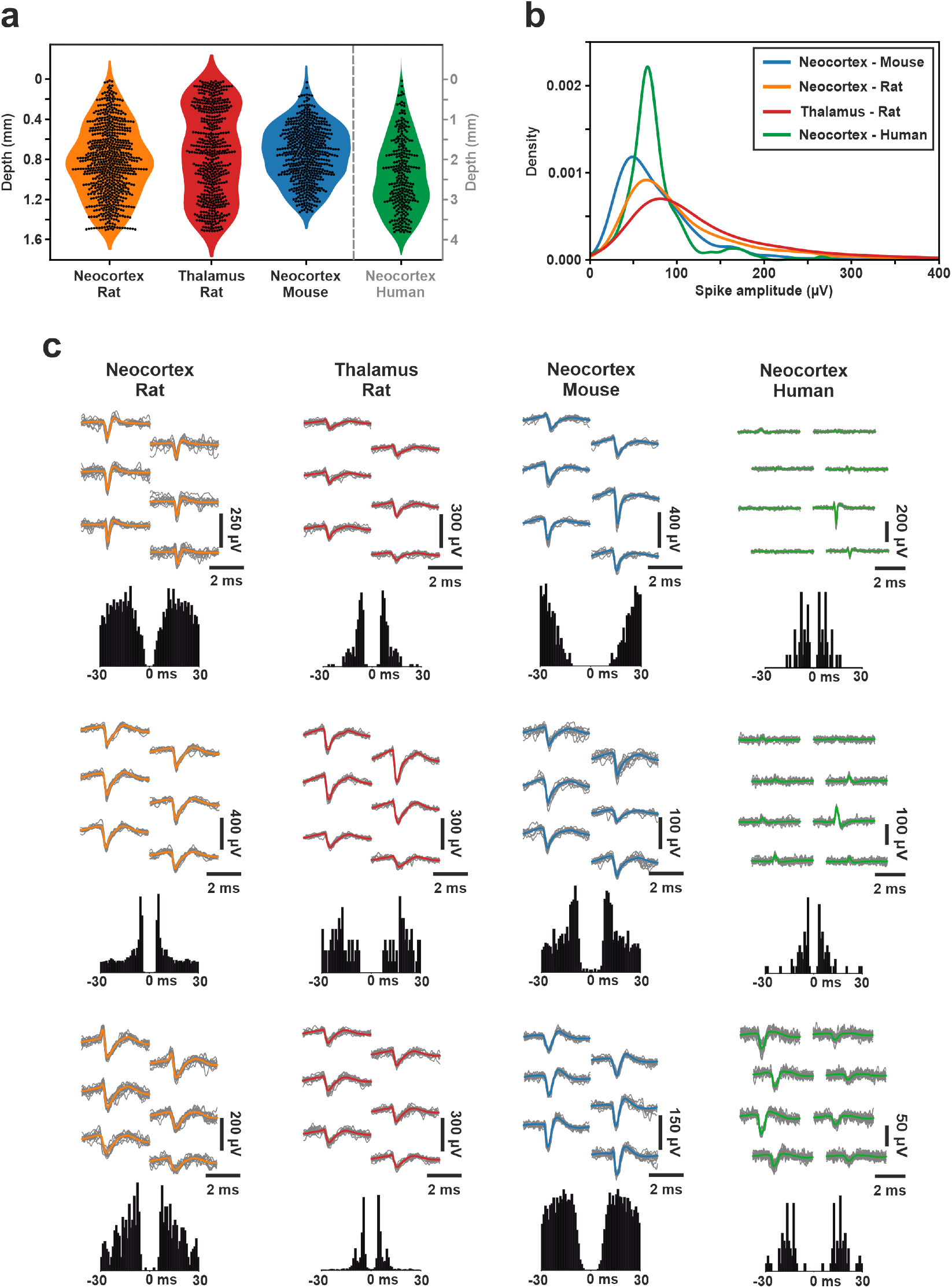
Single unit properties of high-density neuronal datasets. (**a**) Distribution of single unit depths relative to the topmost microelectrode position on the probe. The y-axis corresponding to the human dataset (green) can be found on the right side. (**b**) Distribution of the amplitudes of single unit spike waveforms in each dataset. (**c**) Multichannel spike waveforms and autocorrelograms of three representative single units from each dataset were isolated from the original high-density recordings. For each single unit, individual spike waveforms (n = 20) are colored gray, while the mean spike waveform is overlaid in color. The same color coding is used across all panels.

**Figure 5.**
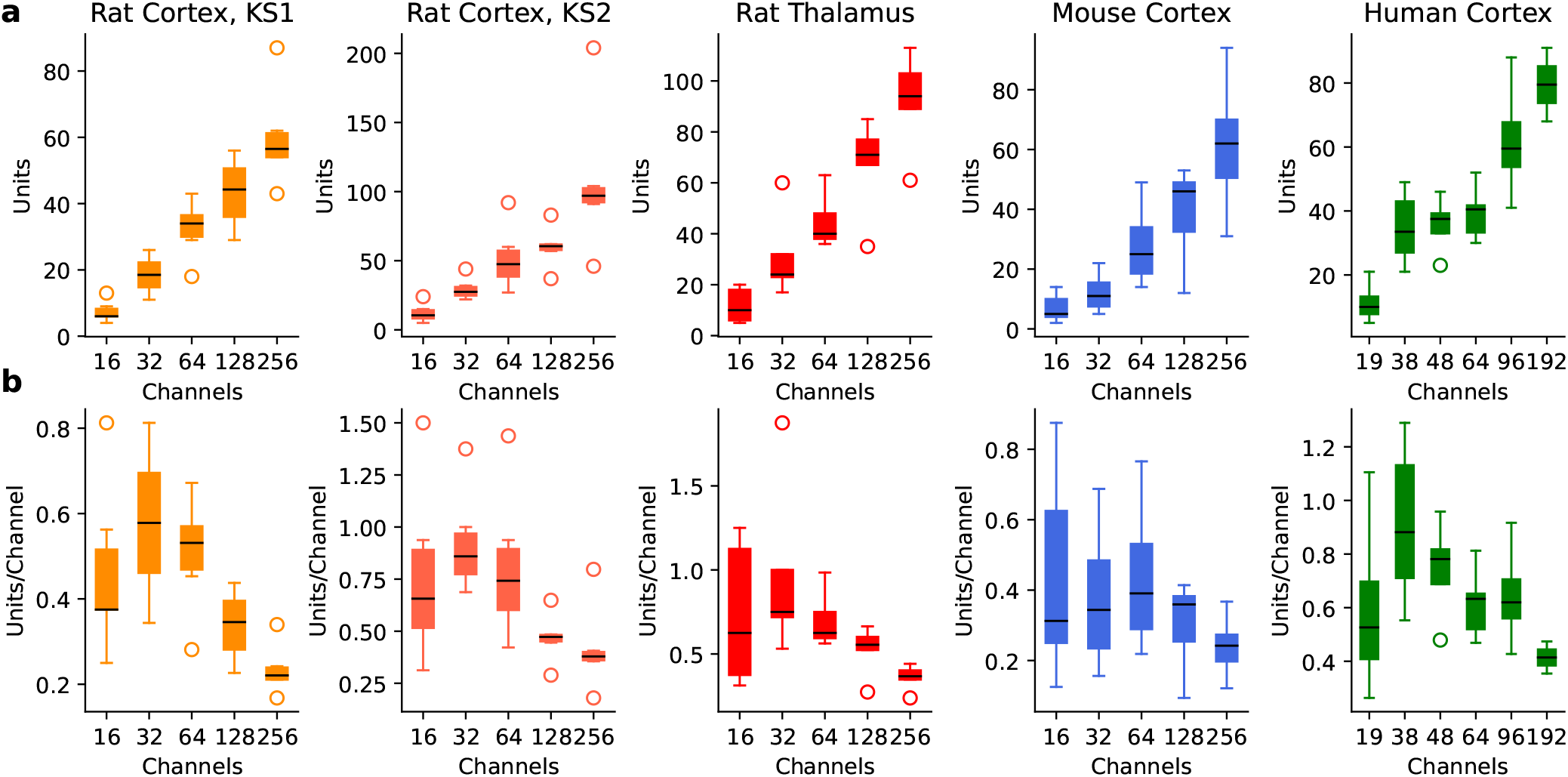
The efficiency of spike sorting using different channel numbers across brain areas, species, and sorting algorithms. **(a)** Median (black lines) and quartiles (colored boxes) of the number of well-isolated single units for various channel counts. **(b)** Efficiency of spike sorting, measured as the unit yield per channel (median and quartiles). While higher channel counts naturally result in a greater unit yield, the efficiency reaches its maximum at a smaller number of electrodes. KS1, Kilosort1; KS2, Kilosort2.

Comparing recordings with different channel numbers (i.e., with varying spatial resolution) revealed a natural trend: more channels (higher spatial resolution or denser sampling) resulted in a higher number of single units (Fig. 5a). However, when assessing spike sorting efficiency — by dividing the unit yield by the number of channels used — we found that the increase in well-isolated single units was not proportional to the increase in channel count (Fig. 5b). The highest channel counts yielded the lowest efficiencies in our comparisons. In all examined cases, the non-monotonic relationship between spike sorting efficiency and channel number suggests the existence of an optimal electrode density for each examined brain area, which may also vary across species.

Since different types of silicon probes with different electrode configurations were used in rodent and human experiments, similar (or the same) channel counts correspond to different electrode densities for the two datasets. To enable a meaningful comparison between measurements, electrode efficiency is presented as a function of inter-electrode distance (Fig. 6). Furthermore, the dual observer model was fitted to the median of spike sorting results (i.e., unit yield per channel; see Methods), and the resulting theoretical data is shown alongside individual measurements and their median (Fig. 6). The theoretical curves of the dual observer model closely match the median of the experimental data in all cases, although the data corresponding to recordings from the rat thalamus exhibits slight deviations from the theoretical curve. The fitted parameters of the dual observer model for all datasets are presented in Table 3.

**Table 3.**
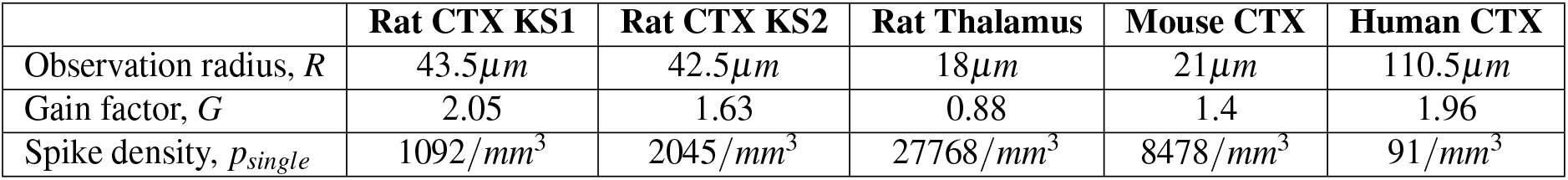
Fitted model parameters. CTX, Neocortex; KS1, Kilosort1; KS2, Kilosort2.

|  | Rat CTX KS1 | Rat CTX KS2 | Rat Thalamus | Mouse CTX | Human CTX |
| --- | --- | --- | --- | --- | --- |
| Observation radius, $R$ | $43.5\mu m$ | $42.5\mu m$ | $18\mu m$ | $21\mu m$ | $110.5\mu m$ |
| Gain factor, $G$ | 2.05 | 1.63 | 0.88 | 1.4 | 1.96 |
| Spike density, $p_{single}$ | $1092/mm^3$ | $2045/mm^3$ | $27768/mm^3$ | $8478/mm^3$ | $91/mm^3$ |

**Figure 6.**
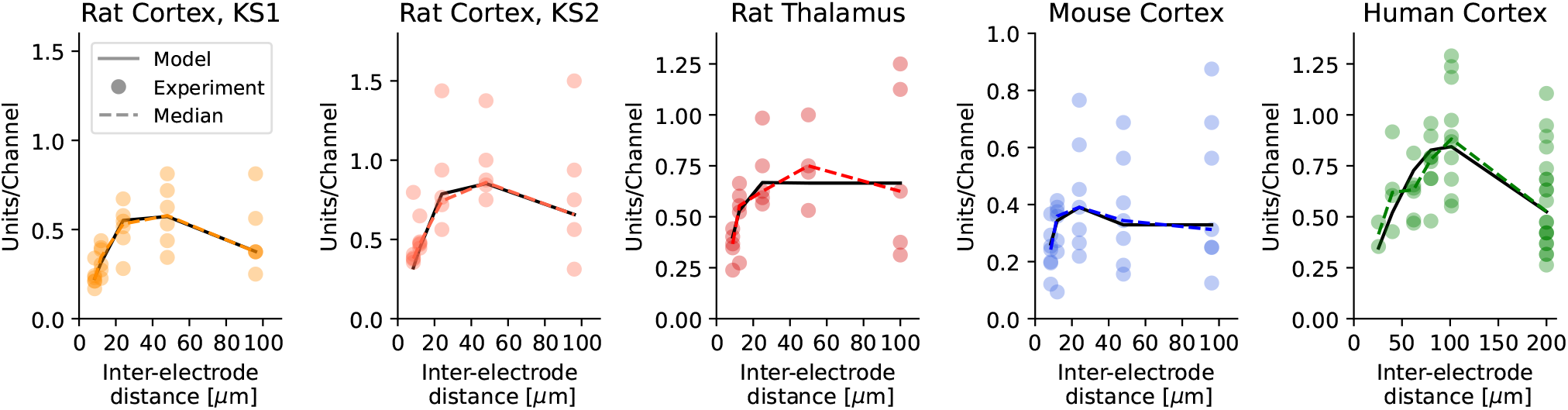
The efficiency of spike sorting as a function of inter-electrode distance across brain areas, species, and sorting algorithms. Colored circles represent units-per-channel results for individual recordings, while colored dashed lines indicate the medians of the measurements. Solid black lines show the best-fitting model. The highest electrode density, corresponding to the smallest inter-electrode distance, was not the most efficient in any of the cases. Instead, an optimal inter-electrode distance was observed across all analyzed recordings. KS1, Kilosort1; KS2, Kilosort2.

The distance dependence of spike sorting efficiency was remarkably similar for both versions of the Kilosort spike sorting algorithm (KS1 and KS2), with one key difference: KS2 identified approximately twice as many units as KS1 (ratio of efficiency *N*_*KS*2_*/N*_*KS*1_ = 1.97 ± 0.63, *mean* ±*SD* for all subsampled recordings). In both cases, the measured efficiency exhibited clear non-monotonicity, with a maximum at 48*μm*. The theoretically derived optimal inter-electrode distances (*D*_*opt*_) were close to these empirical values: 37*μm* for both KS1 and KS2. Note that these *D*_*opt*_ values are given for *M* = 32 electrodes. However, Eq. 4 allows for the calculation of *D*_*opt*_ for any electrode count.

The observed distance dependence of electrode efficiency, characterized by a clear maximum, is a hallmark of the strong cooperative effect between electrodes (Fig. 6). The estimated gain factors were well above 1 in all cases, except for the rat thalamus dataset (Table 4). These findings suggest that a well-defined inter-electrode distance can be determined for these cases. The gain factor calculated for the rat thalamus (G= 0.88) also indicates the presence of a cooperative effect between electrodes. However, this effect is not strong enough to yield a significantly higher number of units to compensate for the volume loss caused by the electrode intersections. From a design perspective, *G* ≤ 1 implies that all electrode configurations where *D > R* have a similar efficiency (compare to Fig. 1 b; blue lines). Within this regime, other factors can determine the optimal configuration. As in most experiments, the size of the tissue to be recorded is limited — for example, the thickness of the cortex constrains the length of the probe shank. Therefore, the optimal electrode configuration is most likely the minimal *D* = *R* distance, as it allows the placement of a higher number of electrodes within the limited volume. The observation radius for the thalamic data was *R* = 18*μm*, thus, all electrode configurations where *D* ≥ 18*μm* have a similar spike sorting efficiency, fitting well to the experimental result, where the maximal unit/channel yield was observed at a spatial resolution of 48*μm* (Fig. 6).

**Table 4.**
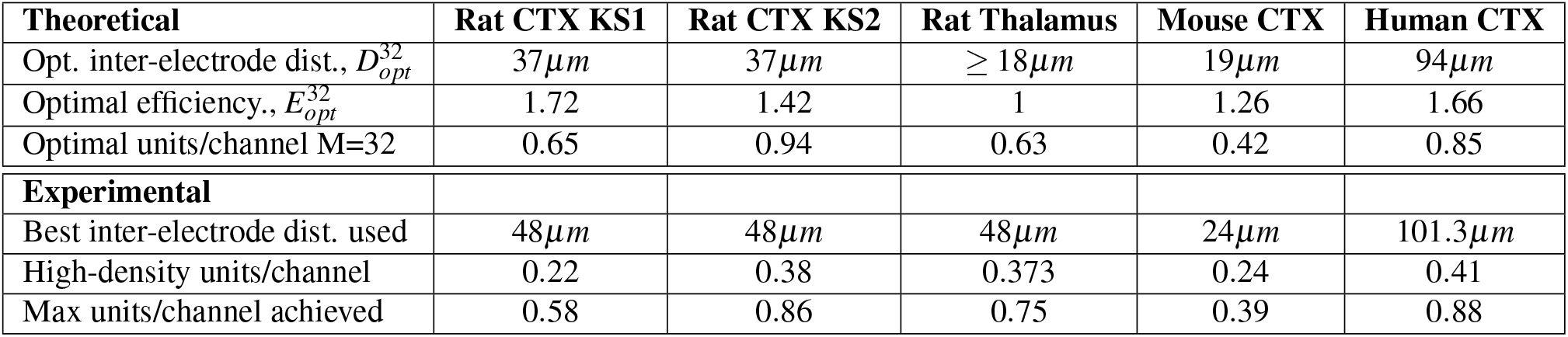
The determined optimal electrode parameters (theoretical results) compared to experimentally achieved values. The optimal inter-electrode distances for a 32-channel linear probe 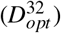, derived from the fitted model parameters, are presented and compared to the inter-electrode distances that resulted in the highest units/channel yield based on in vivo recordings (best inter-electrode distance used). The theoretically estimated and experimentally achieved maximal unit yields (optimal units/channel vs. max units/channel achieved), along with the experimentally best and theoretically optimal inter-electrode distances, were closely aligned. In the case of the rat cortex, theoretical estimates suggest room for further improvement. However, for the rat thalamus, mouse cortex, and human cortex, the downsampled recordings have already reached the theoretical maximum. A comparison of the theoretically achievable optimal units/channel values with the units/channel yield corresponding to the highest electrode densities (i.e., the lowest inter-electrode distance; high-density units/channel) shows that the optimal values are 1.7-3 times higher than the values determined experimentally. CTX: Neocortex; KS1: Kilosort1; KS2: Kilosort2.

The fitted model reproduced fine details of the measured electrode efficiency functions, such as the skewed peaks for all cortical datasets (Fig. 6). The case of the mouse cortex is particularly interesting as it exhibits a clear maximum but becomes flat for larger inter-electrode distances. This behavior exactly matches the distance dependency described by the dual observer model with *G >* 1 (Fig. 1 b, orange and red lines). Above *D >* 2*R* (which is 42*μm* in this case), all the intersections between the observation spheres are eliminated, and the observations of the electrodes become independent. As a result, the number of observed single units no longer depends on the inter-electrode distance in this regime.

Our theoretical results determined for a linear probe with 32 electrodes, summarized in Table 4, lead to several important conclusions. First, there are clear species-specific differences in the optimal inter-electrode distance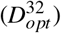. The optimal spacing in the neocortex varied considerably across the three investigated species. The smallest inter-electrode distance was found for mice (19*μm*), while the optimal spacing was nearly twice as large in rats (37*μm*), and approximately five times greater in humans (94*μm*). Second, a brain region-specific difference was observed in rats. The optimal distance between electrodes was almost two times larger in the neocortex (37*μm*) compared to the thalamus (≥ 18*μm*). Third, the theoretical estimation of optimal inter-electrode distances closely matched those found to be most effective experimentally (Table 4). However, in some cases, particularly in rat cortical recordings, there was room for further optimization. Interestingly, the optimal spacing between electrodes was the same for the two versions of the spike sorting algorithm (37*μm*) applied to the rat cortical data. These findings highlight the importance of tailoring electrode configurations to specific experimental goals and brain regions.

The optimal electrode efficiency metric (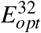 in Table 4) quantifies the improvement of an optimized electrode arrangement compared to widely spaced, independent electrodes. Our results indicate that an optimal configuration improves efficiency by 26 − 72% over independent electrode placements (E = 1), except in thalamic recordings. More importantly, comparing both the experimentally achieved maximal units per channel yields and the theoretically derived optimal units per channel values to those obtained with the highest electrode density configurations reveals a 1.7 − 3 times increase with optimal arrangements (Table 4). This improvement was substantial across all datasets: 70 − 75% increase in unit yield per channel for rat thalamic and mouse cortical recordings, 107% increase in human cortical recordings, and 195%/147% (KS1/KS2) increase in rat cortical recordings.

## Discussion

By fitting our theoretical model to experimental single-unit yield data extracted from high-density recordings and their spatially downsampled versions, we determined optimal inter-electrode distances that maximize the number of well-isolated single units detected with a single electrode. Our findings indicate that the electrode pitch yielding the best results (i.e., the highest unit yield per channel) depends significantly on the investigated brain area and species. For the neocortex, optimal electrode spacing increased with brain size, with nearly a fivefold difference between mice and humans. The thalamus, composed of many small nuclei, exhibited a slightly different pattern and showed the smallest derived electrode spacing. Interestingly, for rodents, the optimal inter-electrode distances were smaller than the electrode pitch of traditional, non-CMOS-based linear silicon probes used in rodent experiments (50*μm* or 100*μm*)^20–22^. Similarly, the best electrode spacing found for the human neocortex (94*μm*) was smaller than the pitch typically used in human intracortical electrode arrays (150*μm* or higher)^23–25^. Our results also demonstrate that spike sorting efficiency corresponding to optimal inter-electrode distances can improve severalfold compared to the electrode configuration with the highest density.

While three parameters might seem like too many degrees of freedom to describe a measurement consisting of only 5-6 points, Tóth et al.^18^ demonstrated that these three parameters can be determined through model fitting on similarly subsampled data. Three parameters are the minimum required to describe a general function exhibiting a maximum; at least three parameters, such as height, width, and location, are necessary to describe a peak. However, different models can use different parameters, so the key question is whether the description can be linked to the physiological properties of the neural tissue and the engineering properties of invasive brain implants, helping us to design better probes. Moreover, the measurements do contain finer details, such as asymmetry or skewness of the peaks or plateaus, which can help assess whether our model provides an accurate description. Considering all these factors, we conclude that the dual observer model accurately described the experimental data, capturing many finer details. Furthermore, the parameters determined through fitting were realistic in their values.

Numerical fitting of the model to the experimental datasets helped us determine the model parameters, which allowed for the calculation of the optimal inter-electrode distance for any number of electrodes. This calculated distance was close to the empirically observed maximum of the unit yield per channel, although it did not match exactly. This discrepancy is because the model fitting and the subsequent calculations take into account the entire shape of the spike sorting efficiency as a function of inter-electrode distances, not just the maximal points.

It is also important to acknowledge the main limitations of our study. First, we assumed that the only limiting factor is the number of channels available for use, and thus we determined the optimal arrangement for fixed electrode counts. However, in real-world scenarios, additional limiting factors may arise. The physical size of the investigated brain area is certainly one of these limiting factors, as it could restrict the number of electrodes that can be placed given a specific inter-electrode distance. It is clear that such a limiting factor would alter the optimal arrangement.

Second, we determined the optimal electrode pitch in only two brain regions: the neocortex and the thalamus. The considerable difference in the optimal spacing of microelectrodes between these regions (37*μm* vs. 18*μm*) suggests that further investigation into other brain areas is needed. Moreover, the unique anatomical structure of various areas (e.g., the layered organization of the hippocampus and the neocortex) suggests that the optimal distance between electrodes may vary even within a single brain region. In general, however, we may assume that areas with higher neuron densities (or smaller neurons) require a higher electrode density, whereas regions with fewer or larger neurons allow for greater spacing between electrodes. Our results also support this notion across species. For instance, previous experimental studies have shown that the average density of neurons in the neocortex is approximately two times lower in rats than in mice, with neurons in rats generally being larger^26–28^. In agreement with this, the optimal inter-electrode distance estimated for the neocortex was about twice as large in rats as in mice (37*μm* vs. 19*μm*). Furthermore, the largest inter-electrode distance among the four datasets was found in the human data (94*μm*), aligning with the lower neuronal density in the human neocortex compared to rodents.

Third, most of the studied neuronal datasets were recorded under anesthesia, except for one human recording from an awake patient. Neuronal activity patterns differ substantially between awake and anesthetized states, not only in temporal firing dynamics but also in the spatial distribution of active neurons^29–33^. Consequently, the optimal inter-electrode distance may also depend on the brain state, although we expect this effect to be relatively small.

Fourth, the algorithm used for spike sorting (Kilosort) may also slightly affect the results^34,35^. Although Kilosort is one of the most widely used and reliable spike sorting algorithms, differences in the single unit yield could arise when compared to other sorters. To mitigate these potential differences, we applied various quality metrics to exclude low-quality units^36,37^. Additionally, the similar optimal inter-electrode distance (37*μm*) found for the two versions of Kilosort suggests that the effect of spike sorters is minimal.

Our findings may aid in developing methods for selecting an optimal subset of electrodes on high-density active (CMOS) probes, or on high-density, high-channel-count multielectrode arrays commonly used in in vitro studies (e.g., to record spiking activity in brain slices or neuronal cultures)^1,3,38,39^. Selecting electrodes at optimal distances could reduce data transmission rates in wireless BCI applications or animal research using chronically implanted probes while maintaining a high single-unit yield per electrode. This approach could also decrease the volume of recorded data in animal experiments; for example, in chronic studies where neuronal activity is continuously recorded with high-density probes for several weeks or months. Species- and region-specific optimal inter-electrode distances could be integrated into the electrode selection software or estimated using short snippets of acquired neuronal activity prior to actual recordings. While not necessarily superior to adaptive electrode selection strategies, this approach may offer a faster, computationally less demanding alternative to scanning each channel and computing metrics (e.g., signal-to-noise ratio) to assess the quality of each channel. Although automated adaptive electrode selection methods might provide higher unit yields, these approaches may not be feasible in certain recording scenarios, such as acute human Neuropixels recordings, where only a few minutes are available to obtain valuable high-density data^8^. In such cases, selecting electrodes at optimal distances determined for the human neocortex could offer a good trade-off between single unit yield and selection time.

In addition to benefiting current neurotechnologies, determining optimal electrode spacings might also aid the design of next-generation brain implants. While CMOS-based high-density active probes are becoming more and more widespread, traditional passive (non-CMOS) silicon probes with lower electrode counts are still widely used in many electrophysiology laboratories^40^. Designing new electrode configurations based on optimal electrode distances could make these passive probes more appealing to researchers, as they produce a more manageable amount of data compared to high-channel-count active probes. In applications where maximizing unit yield is of primary importance but the electrode counts are limited, increasing the spatial resolution is not the most efficient approach. Instead, a given number of electrodes can be utilized most efficiently by optimizing inter-electrode distances to match the specific species, brain area, and sorting algorithm. In the cases investigated, the optimal inter-electrode distances were greater than those achieved by high-density electrodes. Therefore, applying a larger electrode spacing, possibly with multi-shank electrode systems, could result in higher unit counts compared to high-density electrodes.

In this work, we examined only regular (linear) electrode configurations with equidistant spacing. This regular sampling of neural tissue is advantageous when local field potentials (LFPs) are of interest. However, if spiking activity is the primary focus, non-regular patterns should also be considered. In a recent modeling study, Hassan et al.^41^ demonstrated that grouping electrodes into squares of four could be more advantageous than the regular 2D square grid configuration. Although our earlier results^18^ showed that the hexagonal configuration outperforms the square layout among 2D electrode configurations, both theoretical and experimental findings here emphasize that the spacing between microelectrodes plays a critical role in spike sorting efficiency. Therefore, all non-regular grid patterns should be examined to determine whether the observed differences are due to spacing variations or differences in the pattern itself. The application of non-regular grids is a promising direction, and theoretical analysis based on the dual observer model can help clarify this question and guide the identification of the optimal configuration.

In conclusion, we believe that our work represents a step toward a more general optimization framework, where the optimization process could consider additional limiting factors. Beyond the number of electrodes and the size of the brain area, these factors may include tissue damage, the anisotropy of the neuronal electric field, or the direction of electrode implantation.

## Materials and methods

### High-density silicon-based neural probe used to collect in vivo rodent electrophysiological recordings

Neuronal activity in rodents was collected using a single-shank, high-density, silicon-based probe, as described in detail in Fiath et al. (2019)^15^. In short, the device was fabricated on 200-mm-diameter silicon wafers using a commercial 0.13-*μm* CMOS process with a three-metal-layer (AlCu) back-end-of-line. The steps of the fabrication process are described in Fiath et al. (2018)^42^. The implantable shank of the probe is 8 mm long with a cross-sectional area of 100*μm* (width) × 50*μm* (thickness). The shank ends with a chisel-shaped tip, which includes a 300-*μm*-long tapered section. The probes feature a two-dimensional high-aspect-ratio array of 256 square-shaped titanium nitride (TiN) microelectrodes, each with a side length of 5*μm* (Fig. 3a). The microelectrodes cover a total area of 11 × 1535*μm*^2^ and are arranged in two columns (in a zigzag pattern), with a 1*μm* inter-column distance. Each column contains 128 equidistantly spaced electrodes, with a vertical inter-electrode pitch of 12*μm*, providing a high-density spatial sampling of the activity of laminar brain structures. The average impedance magnitude of the TiN microelectrodes was ∼670 kΩ at 1 kHz^15^. Only a small percentage (∼3.5%) of the microelectrodes were defective on the two probes used for recordings (9.06 ± 1.76 nonfunctional electrodes/probe; range: 7-12 electrodes). Each probe was wire-bonded to a four-wing flexible printed circuit board (PCB), which consists of four interface sections for zero insertion force (ZIF) connectors.

### Animal surgery

All in vivo experiments were performed in compliance with the EC Council Directive of September 22, 2010 (2010/63/EU). All procedures were reviewed and approved by the Animal Care Committee of the HUN-REN Research Centre for Natural Sciences and the National Food Chain Safety Office of Hungary (license number: PEI/001/2290-11/2015). In the acute experiments (Table 1), neural recordings were obtained from the neocortex (n = 6 recordings) and thalamus (n = 5) of adult Wistar rats (n = 6; weight: 370.00 ± 140.14g, mean ± SD; n = 5 female; Supplementary Table 1), and from the neocortex (n = 7) of adult C57BL/6J mice (n = 7; weight: 25.73 ± 7.51g; n = 5 female; Supplementary Table 2). All experiments were performed during the subjective nighttime state of the animals.

The surgical and recording procedures were conducted as previously described^15,20,42^. Briefly, anesthesia was induced via intramuscular injection of ketamine (75 mg/kg body weight for rats, 100 mg/kg for mice) and xylazine (10 mg/kg body weight for both rats and mice). Supplementary doses of ketamine/xylazine (1 – 2 injections/hour) were administered as needed to maintain the depth of anesthesia during surgery and recordings. Once surgical anesthesia was achieved, the animals were placed in a stereotaxic frame (David Kopf Instruments, Tujunga, CA, USA). Body temperature was maintained with a homeothermic heating pad connected to a temperature controller (Supertech, Pécs, Hungary). A small craniotomy (3 × 3 mm^2^) was drilled over the brain area of interest. Then, using a 34-gauge needle, the dura mater was pierced to minimize brain dimpling during probe insertion. For rats, the targeted cortical brain areas were the trunk/hindlimb region of the primary somatosensory cortex (S1Tr/S1HL, 4 recordings from 4 rats) and the parietal association cortex (PtA, 2 recordings from 2 rats). Thalamic recordings were obtained from thalamic nuclei located ventrally to these cortical regions (5 recordings from 3 rats). For mice, the probe was inserted into S1Tr (7 recordings from 7 mice). The coordinates of probe insertion and histologically confirmed brain areas are detailed in Supplementary Tables 1 and 2.

For post-mortem histological verification of the recording location^43^, the backside of the silicon probe shank was coated with red-fluorescent dye (DiI, D-282, 10% in ethanol, Thermo Fischer Scientific, Waltham, MA, USA) before penetration. Then, the neural probe was mounted on a motorized stereotaxic micromanipulator (Robot Stereotaxic, Neurostar GmbH, Tübingen, Germany) and driven into the brain tissue at a slow insertion speed (∼ 2*μm*/sec), perpendicular to the cortical surface. To increase the number of recordings, in a subset of rats, multiple probe insertions were performed using the same silicon probe or an intact probe of the same type (see Supplementary Table 1 for details). A minimum distance of 500μm was kept between insertion sites to avoid recording from the proximity of brain tissue damaged by one of the previous penetrations. During the piercing of the dura, as well as during probe insertion, care was taken to avoid large blood vessels located on the brain surface. A room temperature physiological saline solution was applied to the cavity of the craniotomy to prevent dehydration of the brain tissue. A stainless steel needle inserted in the nuchal muscle of the animals served as the external reference electrode during the recordings.

### Acquisition of acute rodent electrophysiological recordings

Spontaneously occurring cortical and thalamic activity was recorded on 256 channels using an Intan RHD-2000 electrophysiological recording system comprising two 128-channel amplifiers (Intan Technologies, Los Angeles, CA, USA) as described in Fiath et al. (2019)^15^. The probe was connected to the amplifiers via a ZIF-to-Molex adapter PCB. Wideband signals (0.1–7500 Hz) were recorded at a sampling rate of 20 kHz/channel with a resolution of 16 bits. Data files containing continuous recordings were saved to hard drives for offline data analysis. Typically, 2–3 hours of neural data were collected from each animal. The main details of the high-density recordings used in this study are provided in Table 1.

### Human Neuropixels recordings

Human cortical Neuropixels recordings^8^ were obtained from the Dryad research data repository^19^. Data from Patient01 (Pt01) and Patient02 (Pt02) were analyzed in this study, both recorded from the dorsolateral prefrontal cortex. The recording from Patient03 contained very low spiking activity; therefore, it was excluded from the analysis. Pt01 was under general anesthesia during deep brain stimulation surgery, while Pt02 was awake at the time of recording. The Neuropixels high-density silicon probe used to collect the human data contains 960 low-impedance TiN microelectrodes, of which 384 can be selected for simultaneous recording^1^. The human variant of the probe has a 10-mm long shank with 70*μm* x 100*μm* cross-section (NP1.0-S^8^). The square-shaped microelectrodes (12*μm* x 12 *μm*) are arranged in a checkerboard pattern with four columns and 480 rows (Fig. 3b). The gap between the columns is 4*μm*, while the gap between rows is 8*μm* (corresponding to 16*μm* horizontal and 20*μm* vertical inter-electrode distances, respectively). Raw recordings were realigned to adjust for vertical tissue movement due to brain pulsations using the automatic Decentralized Registration of Electrophysiological Data (DREDge) algorithm^44^. Neuropixels recordings in the action potential band (AP, band-pass filtered from 0.3 to 10 kHz) were collected using SpikeGLX software (http://billkarsh.github.io/SpikeGLX/) at a sampling rate of 30 kHz/channel. The default electrode map was used for recordings, with the electrodes located on the lower third of the probe (bank0), closest to the probe tip^8^.

### Generation of spatially downsampled electrophysiological recordings

To determine the optimal inter-electrode distances for different brain regions and species, we artificially generated recordings with lower spatial resolutions by selectively removing subsets of channels from the original high-density recordings. These subsampled recordings had the same vertical coverage as the original data. For the 256-channel rodent recordings obtained using the custom laminar silicon probe, the data were spatially downsampled to 128, 64, 32, and 16 channels (Fig. 3a). These linear and equidistantly spaced electrode configurations correspond to inter-electrode distances of 12*μm*, 24*μm*, 48*μm*, and 96*μm*, respectively (Table 2). For the human Neuropixels (384-channel) data, recordings were downsampled to 192, 96, 64, 48, 38 and 19 channels, corresponding to inter-electrode distances of 25.6*μm*, 40*μm*, 62.1*μm*, 80*μm*, 101.3*μm* and 200*μm*, respectively (Fig. 3b; Table 2). Since the single unit yield was higher and the recording quality was better on the channels corresponding to the rightmost two microelectrode columns of the Neuropixels probe (192 channels in total) compared to the two left electrode columns, only the former channels were used to construct the spatially downsampled files. Furthermore, to increase the sample size of the human dataset for a given channel count, we constructed downsampled recordings in each possible electrode configuration by shifting the selected electrodes vertically (Supplementary Fig. 2).

### Spike sorting

Spike sorting was performed on both the original, full-resolution recordings and the spatially downsampled datasets. First, single units were automatically isolated using Kilosort 2.0^34,35^, a MATLAB-based open-source spike sorting software. Following automatic sorting, the Kilosort results were manually reviewed and curated using Phy^45^ (version 2.0), an open-source Python package. For the rodent data, default Kilosort parameters were applied, while for the human data, the parameters were optimized to account for differences in signal characteristics, including sparser neuronal firing rates. Additionally, for cortical recordings obtained from rats, we reused the spike sorting results of our previous study where single unit isolation was performed with Kilosort 1.0, followed by manual curation in Phy^15^.

### Single unit exclusion criteria and quality metrics

For the rodent recordings, a well-isolated single unit was defined as having a clear refractory period and consistent spike waveform shapes. Furthermore, we removed putative single unit clusters with firing rates below 0.1 Hz, as well as duplicated single units and units with very low spike amplitudes (below ∼30*μV*). A small percentage of low-amplitude units were still included in further analysis if other features of the cluster indicated that the spikes were fired by a single neuron.

In contrast to the rodent dataset, the manual curation of the human data was less conservative due to the lower number of good-quality recordings. For example, units with firing rates below 0.1 Hz and lower spike amplitudes were kept to obtain a sufficient unit yield.

For the mouse cortical dataset, single units located outside the cortex (e.g., located in the hippocampus) were excluded from further analysis. The other three datasets contained units only from the investigated brain regions (i.e., thalamus or neocortex). To further improve the reliability of our spike sorting results after manual curation, we removed additional low-quality single units based on various quality metrics. Quality metrics (amplitude cutoff, presence ratio, and interspike interval (ISI) violations) were calculated using SpikeInterface, an open-source Python framework^37^. The amplitude cutoff estimates the false negative rate based on the spike amplitude histogram (units with an amplitude cutoff higher than 0.1 were removed). The presence ratio represents the fraction of time during the recording session when a single unit is active (units with a presence ratio lower than 0.85 were excluded). The ISI violations reflect the rate of refractory period violations (units with ISI violations higher than 2 were discarded). The distribution of these quality metrics for the rodent datasets is shown in Supplementary Figure 3. Approximately 9% of the units were excluded due to these quality metrics. For the human data, the quality metrics-based filtering of units was not applied due to the lower number of isolated units.

### Model fitting

The efficiency of an electrode configuration, according to the dual observer model, can be calculated analytically. First, let us denote the number of successfully identified neurons by *N*. According to the model, *N* is expressed as:

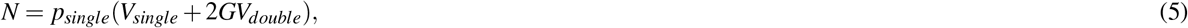

where *V*_*single*_ and *V*_*double*_ are the volumes that are covered by the observation sphere of only one or at least two electrodes, respectively. *p*_*single*_ is the probability density of successful spike sorting if only one electrode observes the electric signal of the neurons, and *G* is the gain factor, expressing the excess probability of successful spike sorting in *V*_*double*_, the volume, where at least two nearby electrodes observe the electric signals of the neurons.

The efficiency of an electrode configuration consisting of *M* electrodes is given relative to *M* independent electrodes as:

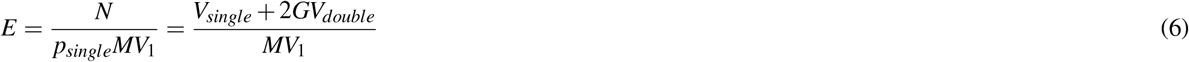

Where *V*_1_ is the volume covered by one single sphere around a microelectrode:

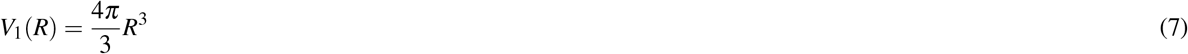

To calculate the electrode efficiency for regular linear electrode configurations, *V*_*single*_, and *V*_*double*_ volumes should be calculated from the intersections of nearby observation spheres as a function of the *R* observation distance and the *D* inter-electrode distance. The details of these calculations can be found in Tóth et al.^18^. Here we summarize the results that were applied to the model fitting. There are three cases:

1. If the electrodes are far away from each other (*D* ≥ 2*R*) then there are no intersections, thus *V*_*single*_ = *MV*_1_ and *V*_*double*_ = 0, thus:

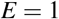

independently of the *G* fgain actor.
2. If 2*R > D* ≥ *R*, then only the first neighbor spheres intersect with each other. The volume of the intersections can be calculated as double spherical caps:

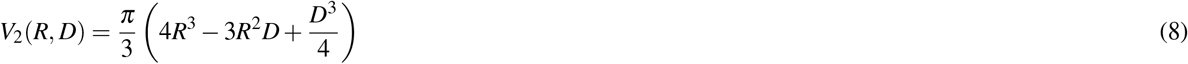

As we have *M* − 1 intersections,

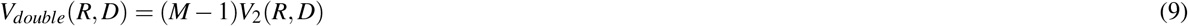

and

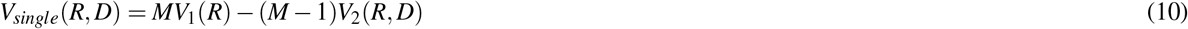

Thus

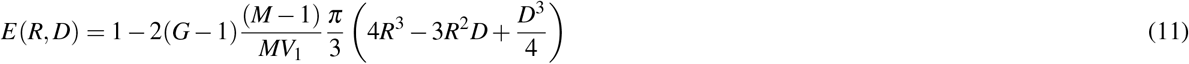
3. If *R*≥ *D*, then *M* − 2 second neighbor intersections also exist and the volume of a second neighbor interaction is *V*_2_(*R*, 2*D*). Thus *V*_*double*_ can be calculated using the inclusion-exclusion principle:

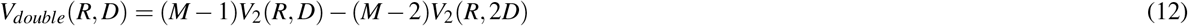

For larger *R*, higher-order intersections may exist, but in a linear electrode arrangement, third-order intersections also create new second-order intersections of the same volumes that cancel each other from the inclusion-exclusion series, thus all the higher-order terms are canceled. Similarly, the total volume can be given with only the first two terms, while the higher-order terms cancel each other:

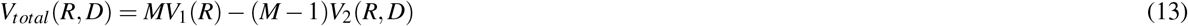

From this:

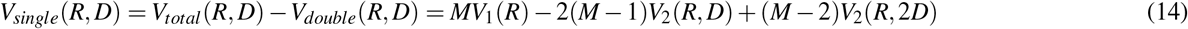

Substituting *V*_1_(*R*) and *V*_2_(*R, D*):

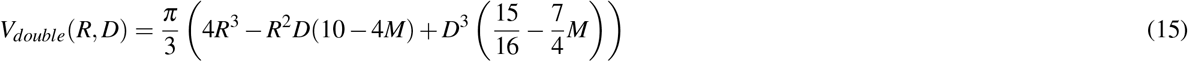

Dividing by the efficiency of the independent electrode model gives us the expression for the efficiency of the linear array as a function of *R, D* and *G*, while the *M* number of electrodes is a parameter:

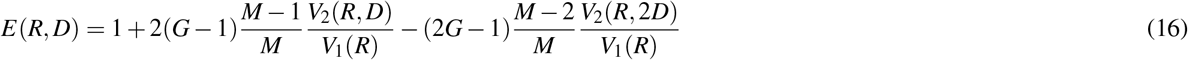

For any of the analyzed datasets, different subsampling rates led to different inter-electrode distances (*D*). Thus, the free parameters of the dual observer model were the *R* observation distance, the *G* gain factor, and the *p*_*Single*_ spike observation density. To search for the optimal model, efficiency patterns were calculated for all 5 (or in the case of human data, 6) different *D* values using the above expressions, where *R* varies in the 1 − 200*μm* interval with 0.5*μm* steps and *G* varies in the 0.5 − 3 interval with 0.01 steps. The calculated efficiency patterns were compared to the median units/channel empirical efficiency values by cosine similarity, which allows amplitude-independent comparison. The most similar pattern resulted in the estimation of the *R*_*opt*_ observation distance and *G*_*opt*_ gain factor, while the *p*_*single*_ probability was estimated from the scalar product of the estimated and calculated efficiency patterns. The estimated *R*_*opt*_ and *G*_*opt*_ values were used to calculate the optimal inter-electrode distances (*D*_*opt*_), according to Eq. 4.

Note that in the case of human recordings, only the even subsamplings followed a strictly linear pattern, while the odd subsamplings exhibited a zig-zag pattern. For our analysis, we applied the linear formulas using the nearest inter-electrode distances computed via the Pythagorean theorem. This approximation remains exact as long as only first-order intersections of observation spheres occur. However, when higher-order intersections start to appear, small second-order deviations may arise. Given the estimated observation distances, these higher-order intersections are expected to appear only in the first or second densest datasets.

## Acknowledgements

The research leading to these results has received funding from the Hungarian Brain Research Program Grant (NAP2022-I-2/2022). R.F. was supported by the Bolyai János Scholarship of the Hungarian Academy of Sciences. Project no. 150574 and 2019-2.1.7-ERA-NET-2021-00023 have been implemented with the support provided by the Ministry of Culture and Innovation of Hungary from the National Research, Development, and Innovation Fund, financed under the STARTING_24 and 2019-2.1.7-ERA-NET funding schemes, respectively. This research was also supported by the Hungarian National Research, Development, and Innovation Office (NKFIH), under grant numbers K135837 (Z.S.), PD143582 (D.M.), and the Hungarian Research Network (HUN-REN) under grant numbers SA-114/2021 (Z.S.) and TECH-2024-20.

## Author contributions

Domokos Meszéna: Investigation, Data Curation, Writing - Original Draft, Writing – Review & Editing. Ward Fadel: Software, Visualization, Writing - Original Draft, Writing – Review & Editing. Robert Tóth: Methodology, Software, Writing – Review & Editing. Tamás Kiss: Writing – Review & Editing, Funding acquisition. Angelique C. Paulk: Data Curation, Writing – Review & Editing. Sydney S. Cash: Writing – Review & Editing, Supervision. Ziv Williams: Writing – Review & Editing, Supervision. Marcell Stippinger: Writing – Review & Editing, Funding acquisition. Lucia Wittner: Investigation, Writing – Review & Editing, Supervision. Richárd Fiáth: Conceptualization, Methodology, Validation, Investigation, Data Curation, Visualization, Writing - Original Draft, Writing – Review & Editing, Project administration, Supervision, Funding acquisition. Zoltán Somogyvári: Conceptualization, Methodology, Software, Formal analysis, Visualization, Writing - Original Draft, Writing – Review & Editing, Supervision, Funding acquisition.

