## Supplementary Information for "Optimal inter-electrode distances for maximizing single unit yield per electrode in neural recordings"

[1]HUN-REN Research Centre for Natural Sciences, Institute of Cognitive Neuroscience and Psychology, Integrative Neuroscience Group, Budapest, H-1117, Hungary
[2]Pázmány Péter Catholic University, Faculty of Information Technology and Bionics, Budapest, H-1083, Hungary
[3]Center for Neurotechnology and Neurorecovery, Department of Neurology, Massachusetts General Hospital, Harvard Medical School, Boston, MA 02114, USA
[4]HUN-REN Wigner Research Center for Physics, Institute for Particle and Nuclear Physics, Department of Computational Sciences, Theoretical Neuroscience and Complex Systems Research Group, Budapest, H-1121, Hungary
[5]University of Oxford, Nuffield Department of Clinical Neurosciences, Oxford, OX1 3TH, UK
[6]Department of Neurosurgery, Massachusetts General Hospital, Harvard Medical School, Boston, MA 02114, USA
[7]Semmelweis University, Faculty of Medicine, Department of Neurosurgery and Neurointervention, Budapest, H-1145, Hungary
[8]Axoncord LLC., Budapest, Hungary
[+]these authors contributed equally to this work

| Recording ID | Animal ID | Target | AP | ML | DV |
|---|---|---|---|---|---|
| 1 | Rat1 | Neocortex (S1HL) | -2.00 | 1.80 | 2.20 |
| 2 | Rat1 | Thalamus (LDVL, AV, VA, VL) | -2.00 | 1.80 | 6.20 |
| 3 | Rat1 | Thalamus (LDVL, Po, VPM) | -2.85 | 2.36 | 6.00 |
| 4 | Rat2 | Neocortex (S1Tr) | -2.70 | 2.51 | 2.20 |
| 5 | Rat2 | Thalamus (LDVL, Po, VPM) | -2.70 | 2.51 | 5.70 |
| 6 | Rat3 | Thalamus (LDVL, Po, VPM) | -2.70 | 2.65 | 5.60 |
| 7 | Rat3 | Thalamus (LDVL, Po, VPM, VPL) | -2.70 | 2.65 | 7.20 |
| 8 | Rat4 | Neocortex (PtA) | -3.45 | 2.42 | 2.10 |
| 9 | Rat5 | Neocortex (S1Tr) | -2.80 | 2.10 | 1.80 |
| 10 | Rat6 | Neocortex (S1Tr) | -2.90 | 2.22 | 1.80 |
| 11 | Rat6 | Neocortex (PtA) | -3.60 | 2.22 | 1.80 |

**Supplementary Table 1.** Stereotaxic coordinates of targeted brain regions in rats. AP, anteroposterior; ML, mediolateral; DV, dorsoventral; S1HL, hindlimb region of the primary somatosensory cortex; S1Tr, trunk region of the primary somatosensory cortex; PtA, parietal association cortex; LDVL, ventrolateral part of the laterodorsal nucleus; Po, posterior nucleus; VPM, ventral posteromedial nucleus; VPL, ventral posterolateral nucleus; AV, anteroventral nucleus; VA, ventral anterior nucleus; VL,ventrolateral nucleus.

| Recording ID | Animal ID | Target | AP | ML | DV |
|---|---|---|---|---|---|
| 1 | Mouse1 | S1Tr | -1.60 | 1.50 | 1.50 |
| 2 | Mouse2 | S1Tr | -1.60 | 1.50 | 1.50 |
| 3 | Mouse3 | S1Tr | -1.60 | 1.57 | 1.80 |
| 4 | Mouse4 | S1Tr | -1.60 | 1.51 | 1.40 |
| 5 | Mouse5 | S1Tr | -1.60 | 1.51 | 1.30 |
| 6 | Mouse6 | S1Tr | -1.50 | 1.59 | 1.30 |
| 7 | Mouse7 | S1Tr | -1.90 | 1.63 | 1.40 |

**Supplementary Table 2.** Stereotaxic coordinates of targeted cortical areas in mice. AP, anteroposterior; ML, mediolateral; DV, dorsoventral; S1Tr, trunk region of the primary somatosensory cortex.

| Dataset | 256 ch | 128 ch | 64 ch | 32 ch | 16 ch |
|---|---|---|---|---|---|
| Rat neocortex | 106.50 ± 52.14 | 60.00 ± 14.64 | 51.67 ± 22.76 | 29.50 ± 7.89 | 12.17 ± 6.75 |
| Rat thalamus | 92.00 ± 19.60 | 67.00 ± 19.13 | 45.00 ± 11.05 | 31.20 ± 16.96 | 11.80 ± 6.87 |
| Mouse neocortex | 61.14 ± 20.10 | 39.14 ± 14.57 | 27.57 ± 12.59 | 12.00 ± 6.22 | 7.00 ± 4.40 |

**Supplementary Table 3.** Average single unit yields across different rodent datasets and recordings (average ± standard deviation). ch, channel.

| **Dataset** | **192 ch**<br>n = 2 | **96 ch**<br>n = 4 | **64 ch**<br>n = 6 | **48 ch**<br>n = 8 | **38 ch**<br>n = 10 | **19 ch**<br>n = 20 |
|---|---|---|---|---|---|---|
| Human neocortex | 79.50 ± 16.26 | 62.00 ± 19.44 | 39.33 ± 8.09 | 36.38 ± 7.01 | 34.40 ± 10.09 | 11.00 ± 4.48 |

**Supplementary Table 4.** Average single unit yields in the human dataset and recordings. The total number of data files generated for each downsampled dataset with different channel numbers is also show (average ± standard deviation)n. ch, channel.

| Dataset | 256 ch | 128 ch | 64 ch | 32 ch | 16 ch | Total |
|---|---|---|---|---|---|---|
| Rat neocortex | 639 | 360 | 310 | 177 | 73 | 1559 |
| Rat thalamus | 460 | 335 | 225 | 156 | 59 | 1235 |
| Mouse neocortex | 428 | 274 | 193 | 84 | 49 | 1028 |

**Supplementary Table 5.** Total number of single units across different rodent datasets and recordings. ch, channel.

| **Dataset** | **192 ch** n = 2 | **96 ch** n = 4 | **64 ch** n = 6 | **48 ch** n = 8 | **38 ch** n = 10 | **19 ch** n = 20 | **Total** |
|---|---|---|---|---|---|---|---|
| Human neocortex | 159 | 248 | 236 | 291 | 344 | 220 | 1278 |

**Supplementary Table 6.** Total number of single units in the human dataset and recordings. The total number of data files generated for each downsampled dataset with different channel numbers is also shown. ch, channel.

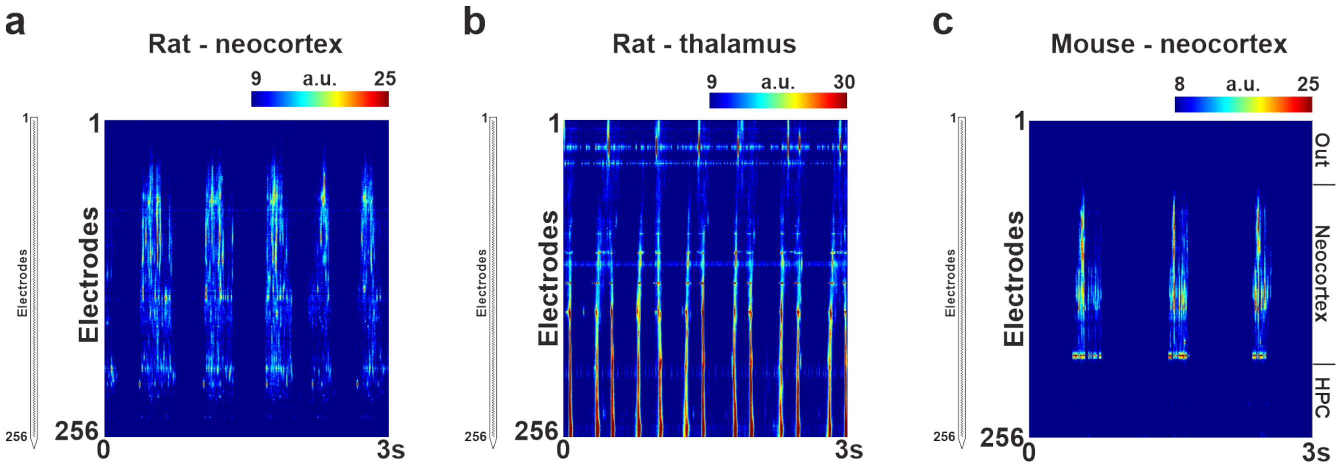


**Supplementary Figure 1.** Representative 3-second-long examples of high-density neuronal recordings (500–5000 Hz frequency band) obtained from the rat neocortex (**a**), rat thalamus (**b**) and mouse neocortex (**c**). On the left, the schematic of the section of probe shank containing the microelectrodes (small black squares) is displayed. On the right, color maps visualizing spiking activity across all channels (i.e., depth profiles) are presented. For these depth profiles, before plotting, data on each channel was rectified, then smoothed with a 50 Hz low-pass filter (third-order Butterworth filter). Warmer colors on the color maps indicate higher spiking activity. Rat and mouse data were acquired under ketamine/xylazine anesthesia. HPC, hippocampus.

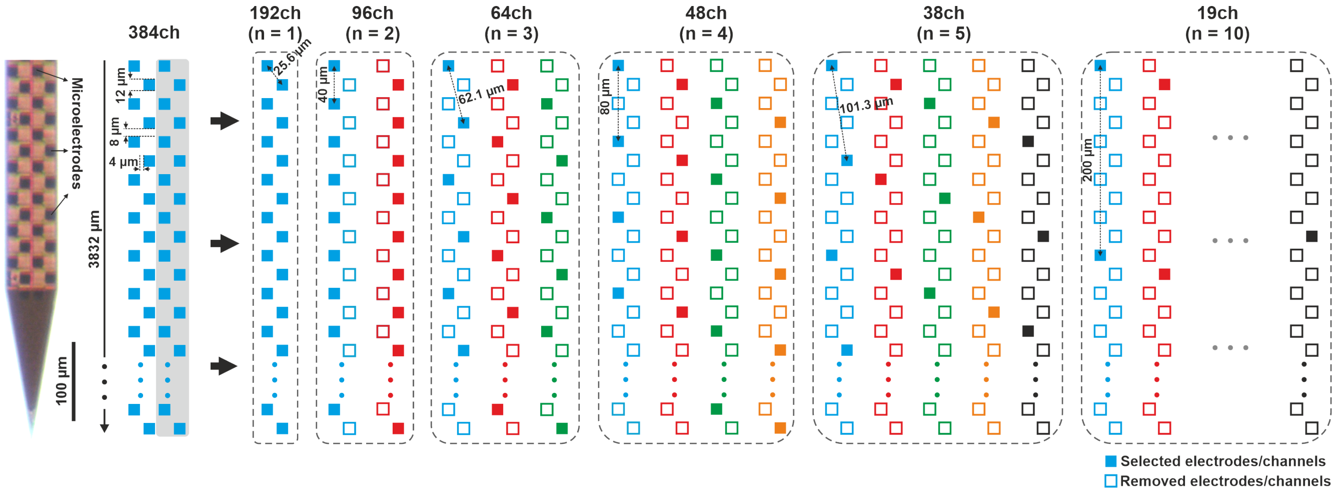


**Supplementary Figure 2.** Spatial downsampling of high-density Neuropixels recordings. Lower channel (ch) count recordings, corresponding to reduced spatial resolutions, were generated from the original 384-channel human cortical recordings (spatial resolution of recordings decreases from left to right). Only data recorded by the rightmost two columns of electrodes (gray shaded area) were used for analysis. To increase the sample size, all possible electrode configurations were generated for each spatial resolution (channel count). Different configurations are indicated with different colors. The size of the microelectrodes, along with the inter-electrode distances for both the original and downsampled recordings, are displayed. A stereomicroscopic image of the tip region of the Neuropixels probe used for human recordings is shown on the left.

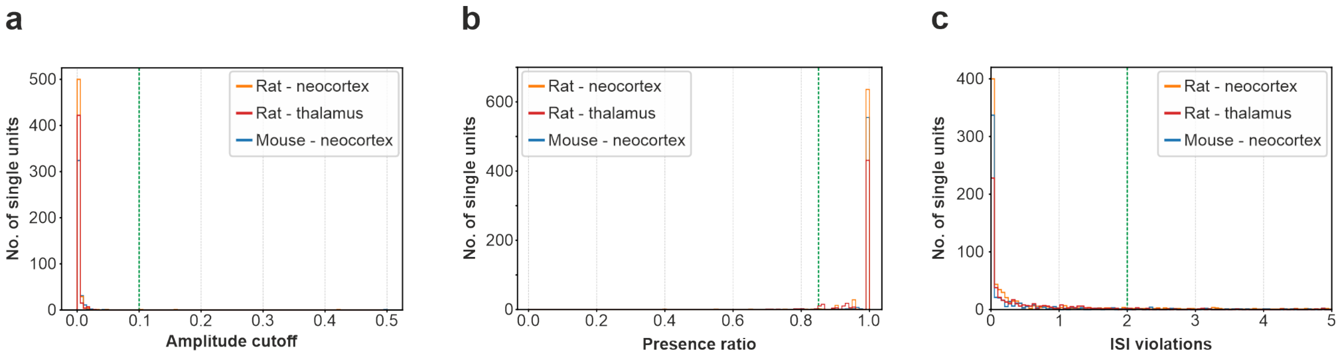


**Supplementary Figure 3.** Distribution of different quality metrics for the rat and mouse datasets. (**a**) Amplitude cutoff, (**b**) presence ratio, (**c**) interspike-interval (ISI) violations. The vertical dashed green lines indicate the thresholds used to exclude low-quality single units.
